# Inferring a qualitative contact rate index of uncertain epidemics

**DOI:** 10.1101/107474

**Authors:** Marco Tulio Angulo, Jorge X. Velasco-Hernandez

**Affiliations:** Institute of Mathematics, Universidad Nacional Autónoma de México (UNAM), Juriquilla 76230, México

**Keywords:** Epidemic prediction, contact rate, qualitative, uncertainty

## Abstract

We will inevitably face new epidemic outbreaks where the mechanisms of transmission are still uncertain, making it difficult to obtain quantitative predictions. Thus we present a novel algorithm that qualitatively predicts the start, relative magnitude and decline of uncertain epidemic outbreaks, requiring to know only a few of its “macroscopic” parameters. The algorithm is based on estimating exactly the time-varying contact rate of a canonical but time-varying Susceptible-Infected-Recovered epidemic model parametrized to the particular outbreak. The algorithm can also be extended to other canonical epidemic models. Even if dynamics of the outbreak deviates significantly from the underlying epidemic model, we show the predictions of the algorithm remain robust. We validated our algorithm using real time-series data of measles, dengue and the current zika outbreak, comparing its performance to existing algorithms that also use a few macroscopic parameters (e.g., those estimating reproductive numbers) and to those using a thorough understanding of the mechanisms of the epidemic outbreak. We show our algorithm can outperform existing algorithms using a few macroscopic parameters, providing an informative qualitative evaluation of the outbreak.

## I. INTRODUCTION

New epidemic outbreaks of influenza, ebola, dengue, zika or other novel diseases are waiting to emerge [1]. For some of these new outbreaks, the mechanisms determining key characteristics such as disease transmission, risk groups, contact rates, vectorial capacities and so on will be uncertain. A thorough understanding of such mechanisms is necessary to obtain quantitative predictions of an outbreak (e.g., its start, magnitude and decline), because they are used by the detailed mathematical models generating the predictions [2–4]. Since these mechanisms can rapidly become very complex due to genetics [5], human mobility [6] and many other factors such as climate change [1], developing accurate quantitative predictions of new outbreaks is a process that can take time—time to collect detailed data of the epidemic outbreak and time to fully understand and mathematically model its mechanisms [7]. For epidemic emergencies or in the unlawful release of infectious agents, time is critical. And thus, before we fully understand the mechanisms of an epidemic outbreak, it is useful to have an early evaluation of the timing and size of the outbreaks—even if the evaluation is just qualitative.

The “macroscopic” dynamics of the majority of past and present epidemic diseases can be reasonable captured by simple canonical epidemic models, such as the Susceptible-Infected-Recovered (SIR) or Susceptible-Exposed-Infected-Recovered (SEIR) ones [8–10]. By contrast to detailed models, these canonical models require only a few “macroscopic” parameters of the outbreak (e.g., birth/mortality and recovery rates of the population). But even with these few parameters, the available data can result in a large uncertainty in their value [10]. This uncertainty might increase further in the case of emergent epidemics, as happened in the recent zika epidemic [11].

Here we propose an algorithm to qualitatively predict the start, relative magnitude and decline of an epidemic outbreak, requiring to know only a few of its “macroscopic” parameters. The algorithm is based on robustly estimating the time-varying <u>contact rate</u>—the fraction of the susceptible population that an average infectious individual successfully infects per day—of a canonical but time-varying SIR (tvSIR) epidemiological model parameterised to the particular outbreak. The contact rate and the related basic/effective reproduction numbers are key quantities for predicting and managing the start, magnitude and end of epidemic outbreaks [8, 12–14]. Hence, a significant number of studies have approached the problem of estimating these quantities from time-series of infection incidence. Most of these efforts focus on estimating constant contact rates and the basic reproductive number [15, 16], with less frequent attempts to estimate time-varying contact rates [17–20]. Some of these estimation methods, such as those in [18, 20], also require few macroscopic parameters of the outbreak. However we show they tend to incorrectly predict, even qualitatively, the start, relative magnitude and decline of an outbreak.

Our algorithm is similar to these last works in that it also relies on incidence time-series data. Indeed, despite being an aggregated measurement of possibly very complex transmission processes—involving immunity, infectiousness, resistance and so on— this is often the first and only available data of an emergent outbreak. We rigorously prove that the proposed algorithm estimates exactly the contact rate in the ideal case when the outbreak dynamics follows exactly a tvSIR model (Section II-A); numerically show its robustness to parametric uncertainties and unmodeled dynamics (Section II-B); and experimentally demonstrate on data of measles, dengue and zika that our algorithm can outperform the qualitative predictions of similar algorithms that only use macroscopic parameters of the outbreak (Section II-C). The algorithm also provides information on how much the actual dynamics of the outbreak differs from its proposed tvSIR model. This allows, for example, detecting if the outbreak is decaying faster than expected by the model. We end the paper offering some concluding remarks and discussing the limitations of the algorithm (Section III).

## II. RESULTS

Assume that incidence time-series data of the epidemic is available, reporting the proportion of the infected population *I*(*t*) at discrete time instants *t* ∈ 𝕋 = {*t*_0_,…,*t_f_*}. Using this data, our qualitative contact rate index takes the form

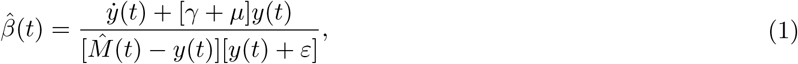

where *M̂* (*t*) is the solution to the differential equation *M̂*̇ (*t*) = *µ* − *µ M̂* (*t*) − γ*y*(*t*) with initial condition *M̂* (*t*_0_) = 1 − (γ/*µ*)*y*(*t*_0_). The continuous-time signal *y*(*t*) in (1) is obtained by interpolating (and filtering, if needed) the data {*I*(*t*),*t* ∈ 𝕋}, and ẏ (t) is an estimate of its derivative at time *t*. Note that (1) is negative when the outbreak develops in a faster time-scale than the one imposed by the parameters of *M̂* (see Section II-B for details). The parameters *µ* > 0, γ > 0 and *ε* > 0 represent the rate of birth/mortality, infected recovering rate, and a small migration term of the outbreak, respectively. These three “macroscopic” parameters depend on the disease and population only, and thus can be adjusted according to previous outbreaks of the same disease, or heuristically adjusted using existing data (see Section II-C).

In what follows we rigorously analyse the performance of (1) when the outbreak dynamics follows an ideal tvSIR model; validate its robustness to deviations of the outbreak dynamics from this model; and analyse and compare its performance using real data.

### A. Theoretical analysis

Consider that the outbreak dynamics follow a canonical tvSIR model [21] with parameters *µ*, γ and *ε*, and a time-varying contact rate *β* (*t*):

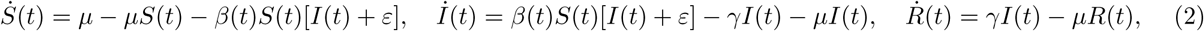

where the population is composed of susceptible *S*, infected *I* and recovered *R* individuals—scaled as proportions of the total population. In this case, using algorithm (1) with *y*(*t*) = *I*(*t*) guarantees that *β̂* (*t*) → *β* (*t*) as *t* →∞ exponentially fast with rate *μ*(*t* − *t*_0_), Fig 1a. In order to prove this claim, let *M* = *S* +*I* and note its time derivative is *Ṁ* = *µ* − *µM − γI*. Hence, using the choice for *M̂* ̇ in (1), we obtain

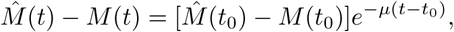

implying that *M̂*(*t*) converges to *M*(*t*) exponentially fast with rate *µ*(*t*— *t*_0_). Consequently, the quantity *Ŝ* (*t*) = *M̂*(*t*) *− y*(*t*)—obtained from data and the numerical solution of (1)— also converges exponentially fast to *S*(*t*), <u>regardless of the particular contact rate</u>, Fig.1b. Borrowing the terminology of system and control theory, *Ŝ*(*t*) and the dynamics for *M̂*(*t*) are an unknown input observer for (2), see e.g. [22] and [23]. This means that, by measuring *y* = *I* only, the *S* variable of the tvSIR model is <u>detectable</u>, implying that it is impossible to estimate the proportion of susceptible population faster than the exponential rate provided by our algorithm. Finally, using *Ŝ*(*t*), we can obtain the contact rate *β* (*t*) using the second equation in (2), yielding the formula for *β* ̂ in (1) and thus proving the claim.

**FIG. 1:**
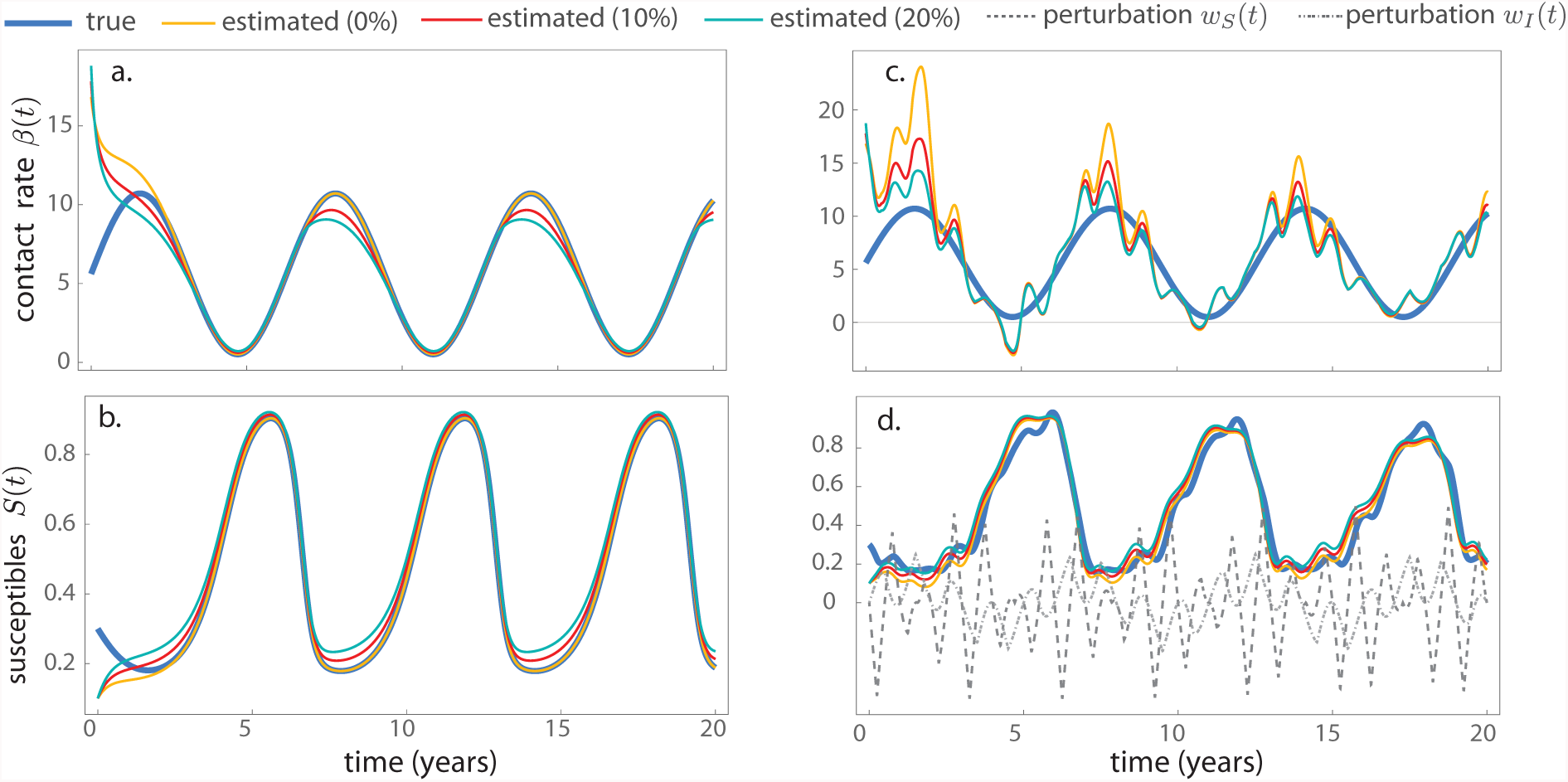
Numerical results. Performance of (1) on the tvSIR model (2) with *γ* = 0.5, *µ* = 1.5, *ε* = 10^−2^ and *β*(*t*) ***=*** 5.1(sin(*t*) + 1.1). **a. and b**. When the proposed algorithm uses exactly the parameters of the tvSIR model, the estimation of *β* (*t*) and *S*(*t*) is exact and exponentially fast. And even when there exists 10% or 20% mismatch between the parameters of the tvSIR model and the ones used by the algorithm (i.e. (1) uses the parameters 1.1*µ*, 0.9*γ*, 1.1*γ* or 1.2*µ*, 0.8*γ*, 1.2*γ*), the proposed algorithm provides a robust estimation. **c. and d.** The robustness of the algorithm allows qualitatively tracking the trend of *β* (*t*) despite the perturbations *w_S_*(*t*) = − 0.51 cos(*t*)|*w*(*t*) and *w_I_*(*t*) = 0.3sin(cos(2*t*))*w*(0.5*t*) acting on the model (here *w*(*t*) is a triangular wave with period 1 varying in [−1, 1]).

By using *β̂* (*t*), we can also construct an index for the effective (or instantaneous) reproductive number:

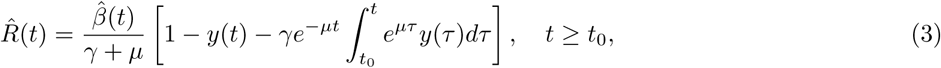

This quantity preserves the usual property of reproductive numbers: the condition *R̂*(*t*) *<* 1 is sufficient for the global asymptotic stability of the infected-free solution *I*(*t*) ≡ 0 of the tvSIR model (2), see Supplementary Information 1. Note that at the start of an epidemic (i.e., when *I* ≈ 0) and assuming a constant contact rate *β*, the formula (3) reduces to the well-known basic reproductive number *R*(0) = *β/*( *γ* + *μ*) for the (time-invariant) SIR model [15]. Additionally, for the experimental results discussed in Section II-C, the approximation *R̂*(*t*) ≈ β̂ (*t*)*/*(*γ* + *μ*) is reasonable, since the magnitude of *y* is of the order 10^−3^.

### B. Numerical validation of its robustness

No outbreak dynamics can be exactly captured by the tvSIR model (2). Thus, in general, we cannot expect to obtain an exact estimation of the contact rate of the outbreak using (1). Instead, the proposed algorithm must qualitatively follow the contact rate of an epidemic outbreak, even if its dynamics significantly deviate from the tvSIR model due to incomplete information on cases, underreporting, demographic and environmental stochastic effects, etc. This requires robustness of the estimation of *β* (*t*) to both changes in the parameters of the tvSIR, and the presence of unmodeled dynamics or perturbations in dynamics of the outbreak.

On one hand, by the continuity of Eq. (1) with respect to its parameters, the estimator *β*(*t*) is robust to changes in the parameters *γ*, *μ* and *ε*, see Fig. 1a-b. This means that a small deviation of these parameters from the “true ones” will only produce a small error in the estimation of the contact rate. Numerical experiments suggest that (1) performs better when the birth/mortality rate *μ* is overestimated and the recovering rate *γ* is underestimated (Fig. S1).

On the other hand, in order to analyse the performance of the proposed indicator in more general scenarios were the dynamics of the outbreak significantly deviates from tvSIR model, we considered the <u>perturbed</u> tvSIR model

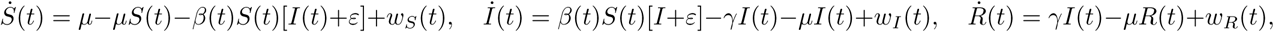

where *w_S_* (*t*), *w_I_* (*t*) and *w_R_*(*t*) are perturbations representing unmodeled mechanisms of infection, demography, vaccination and variability in recovery infection and mortality rates. Even when these perturbations have the same order magnitude as the model (i.e., they have the same order of magnitude as *S* itself), we find that the estimation *Ŝ* (*t*) of *S*(*t*) from (1) remains acceptable and robust to changes in the parameters also (Fig. 1d). In addition, provided that the perturbations occur at a faster time scale than changes in the contact rate, the indicator *β̂*(*t*) follows closely the trend of the true contact rate, and it is also robust to changes in the parameters of the algorithm (Fig. 1c).

Note that due to the presence of these perturbations, the quantity *β̂* (*t*) can be negative. This happens when *I*(*t*) is decreasing faster than the exponential *e*^**−(**γ**+** *μ*)*t*^. Hence, algorithm (1) also measures the discrepancy between the data and the model assumptions. Although this can complicate its interpretation as a true “contact rate”, in Section II-C we argue this is compensated by its better predictive properties compared to other standard contact rate approximations.

### C. Experimental results

We tested the proposed indicator on time-series data of measles in New York, dengue in Rio de Janeiro and the current zika epidemic in Brazil and Colombia (see Methods for details of the data and parameters of the algorithm). We also compared its performance with recent methods for estimating the time-varying reproductive number [14, 18, 20]. In the first two references, the estimation is based on the quantity

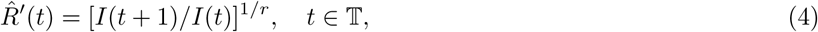

for some exponent *r* (typically equal to one), together with an additional statistical treatment for improving the estimation. In the last reference [14], the authors use the more sophisticated statistical inference method described in [19]. The methods of the first two references require few “macroscopic” parameters of the outbreak, similar to the proposed algorithm. The method in the third reference [20] thoroughly exploits the detailed mechanisms of the zika epidemics, and thus it is expected to be quantitatively and qualitatively better. The results of the comparison are shown in Fig 2.

**FIG. 2:**
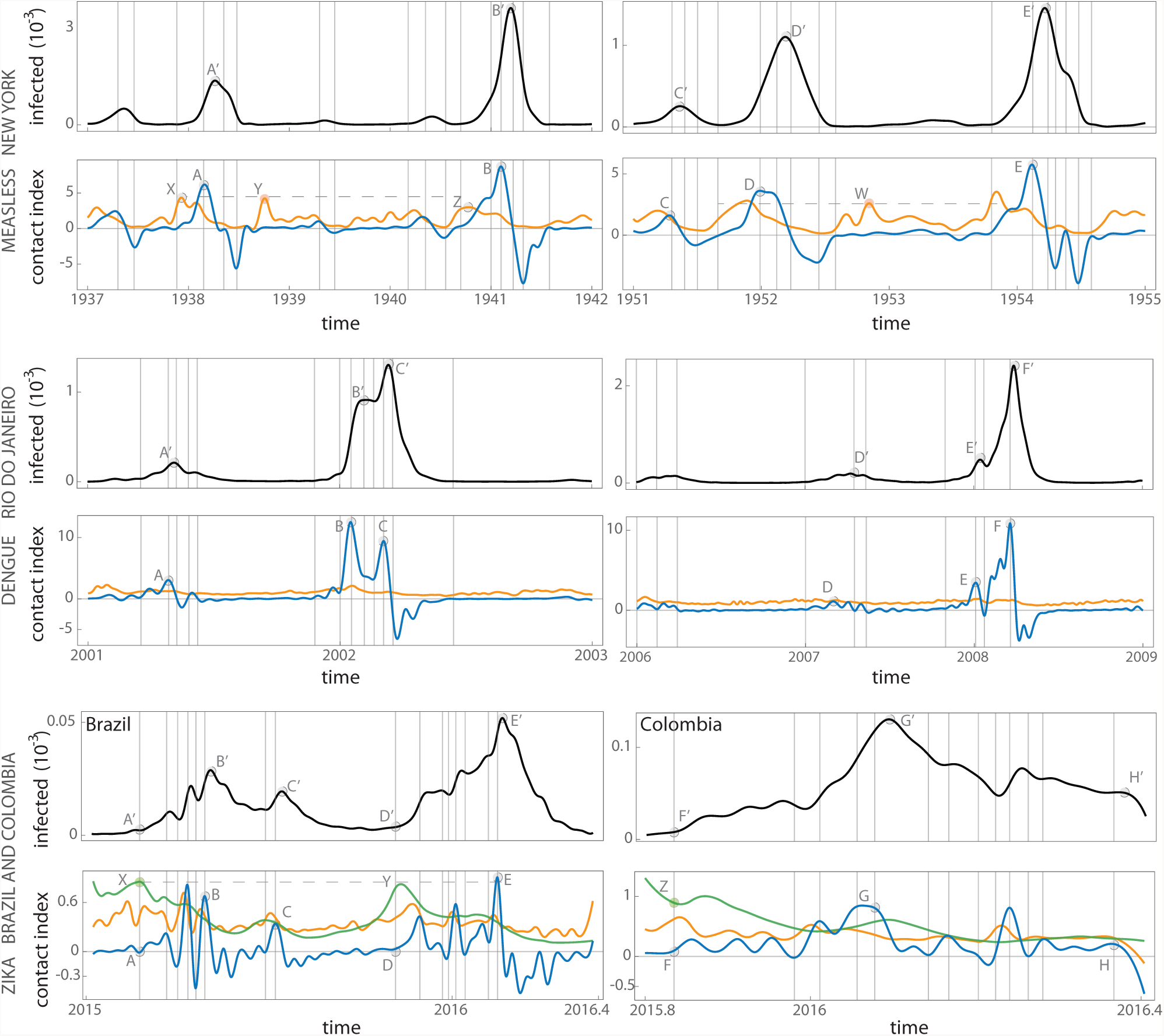
Experimental results on reported cases of measles in New York, dengue in Rio do Janeiro (Brazil) and the current zika epidemic in Brazil and Colombia. Blue: *β̂*(*t*) in (1). Orange: results from the algorithm for estimating the (time-varying) reproductive number *R̂′* (*t*) based in Eq. (4). Green: the (mean) estimation for the time varying reproductive number obtained in [14] for the zika epidemic

In the case of measles—with outbreak dynamics reasonably modeled by a SIR model—we observe that *β̂* (*t*) is able to predict early the maximum of the epidemic outbreaks: points *A* to *E* mark months ahead the maxima observed in the data for outbreaks *A*′ to *E*′. Note that the algorithm (4) predicts earlier than ours those maxima (e.g., the point *X* precede point *A*), but this algorithm also predicts non-existent outbreaks (points *Y* and *W*, for example). Furthermore, the magnitude of *R̂′*(*t*) does not correlate correctly with the magnitude of the outbreak. For instance, point *X* has larger magnitude than point *Z*, but the corresponding outbreaks have the oposite trend: *A*′ is way smaller than *B*′. By contrast, for the proposed algorithm, the larger the magnitude of a peak in *β̂*(*t*) the larger the outbreak is, showing it correctly predicts the relative magnitude of the outbreaks. For instance, *A < B* and *A′ < B′*.

In the case of dengue, we expect the tvSIR model to be a poor approximation of the outbreak dynamics, since it ignores important mechanisms such as serotype diversity and vector dynamics. Nevertheless, our indicator still qualitatively predicts reasonably well the relative magnitude, start and end of the outbreaks; although its predictive power (e.g., how far ahead can the start or end of the outbreak be predicted) decreases compared to the case of measles. In this case, we also find that the algorithm (4) provides no useful information of the outbreak.

Finally, in the case of zika, for the same reasons as with dengue, we also expect that a tvSIR poorly models its mechanisms. Yet, we observe that our indicator can predict weeks ahead the start (e.g., *A* and *A′* or *D* and *D′)* and the decline (e.g., *B* and *B′* or *E* and *E*′) of the epidemic. We note that its predictive power is similar to the estimation obtained using the algorithm [19] (e.g., points *X* and *A* similarly predict the start of the epidemic *A*′), but our indicator still outperforms the other since it gives a better estimation of the relative magnitude of the outbreak. For example, points *X* and *Y* given by the algorithm in [19] suggests that the magnitude of the outbreaks should be very similar in both cases, but in reality *B′* < *E′*. On the other hand, the predicted magnitude given by our indicator satisfies *B* < *E*.

The proposed indicator can also be used to obtain a quantitative prediction of the infected population in a given time interval. To do this, the value of *β̂*(*t*) on a time interval [*t_n_*, *t_n_* + *ε*] is estimated from its values on [*t*_0_, *t_n_*] by using e.g. splines, echo-state neural networks or autoregressive discrete-time linear models. Then, the expected proportion of infected population *Î*(*t*), *t* ∈ [*t_n_,t_n_* + *ε*], can be obtained by simulating the tvSIR model (2) over the interval [*t_n_,t_n_* + *ε*] with initial conditions *Î* (*t_n_*) = *I*(*t_n_*), *Ŝ* (*t_n_*) = *M̂* (*t_n_*) − *I*(*t_n_*) and contact rate *β̂* (*t*). We tested this method to predict the proportion of infected population in the big measles outbreak in New York near the year *t_n_* = 1941.18, and the dengue outbreak in Rio do Janeiro near the year *t_n_* = 2002.18, see Methods for details. Our method provides an adequate prediction for *ε* = 3.6 months in the case of measles, and *ε* = 18 days in the case of dengue, Fig. 3. We can also correctly predict the increase and latter decline of the infected population.

**FIG. 3:**
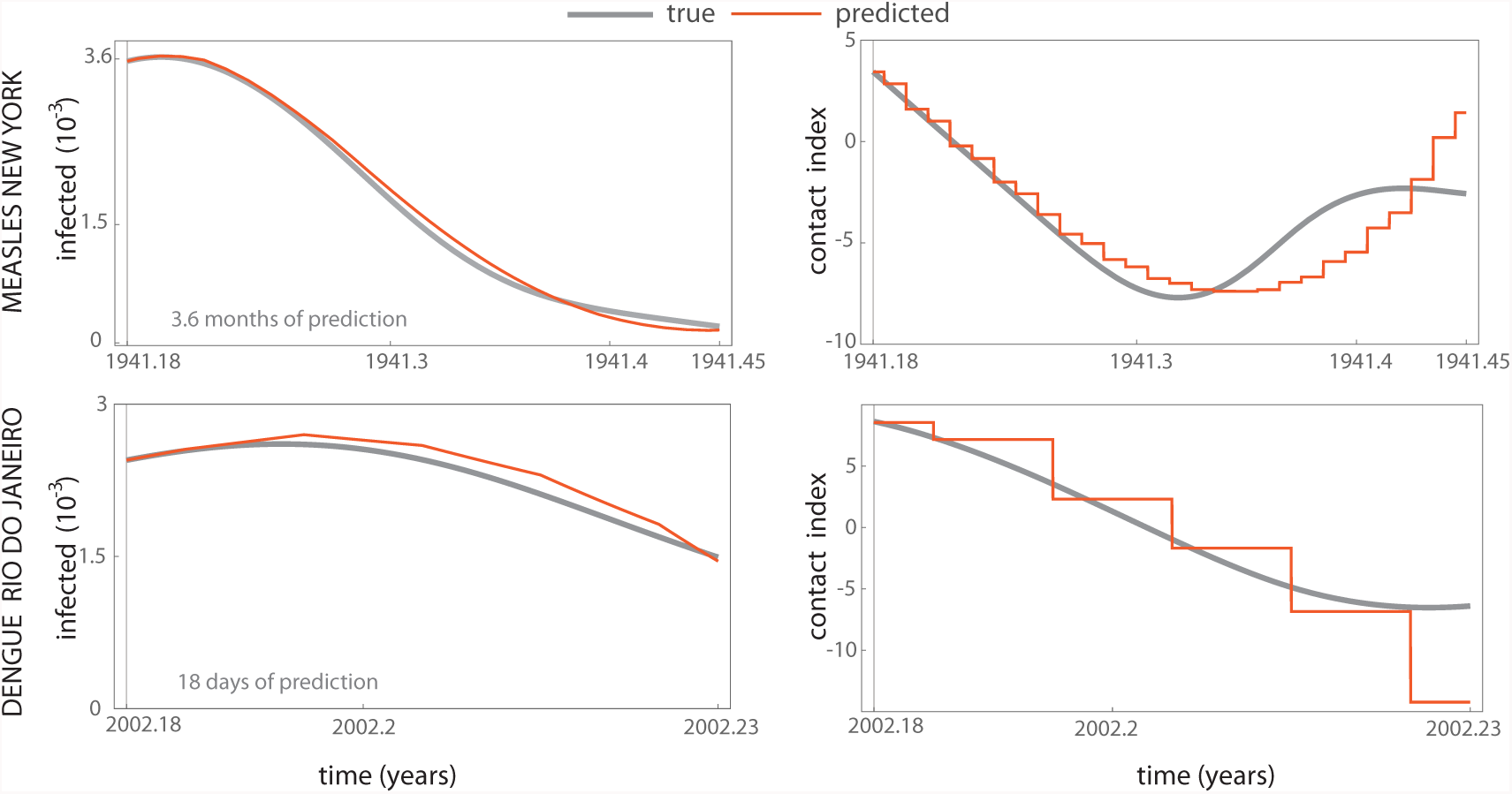
Prediction of proportion of infected population for measles in New York and dengue in Rio do Janeiro. The proposed method provides an acceptable prediction for 22 days in measles, and 2 days in dengue.

## III. DISCUSSION AND CONCLUDING REMARKS

By aiming for a qualitative instead of a quantitative prediction, we have proposed an algorithm that only uses macroscopic parameters of the outbreak, based on an elementary tvSIR epidemiological model. The proposed estimator (1) can be interpreted as an indicator closely associated to the true contact rate. The design of the proposed algorithm borrows ideas from the so-called “unknown input observers”, which have been very effective algorithms for solving estimation problems in engineering [22, 23]. The main property of the tvSIR allowing the design of such observers is its detectability with respect to the infected population. Since other common but more elaborate epidemi-ological models such as SEIR, SIS and so on have this same detectability property, Eq. (1) can be straightforwardly modified to replace the tvSIR model by any of those other models. This would allow incorporating more information of the mechanisms of the epidemic outbreak on the qualitative predictions generated by the proposed algorithm. For example, in the case of dengue, it is reasonable that the model includes state variables for each serotype and the mosquito population. We note that despite the success of observers in solving diverse engineering problems, their application to epidemiological models seems rather unexplored. Indeed, we were only able to locate the recent work [24] that considers the classical SIR model. Note also that our algorithm is similar in spirit to [17], in which the authors statistically fit the parameters and initial conditions of a time-invariant SIR model to time-series data in order to estimate the effective reproductive number.

An intrinsic limitation for applying the proposed algorithm is that the data needs to have enough samples to adequately represent the continuous time evolution of the infected population. Rigorously, according to the Nyquist criterion, this require sampling at more than twice the frequency of the bandwidth of this continuous time signal. Note also that for highly noisy data, the estimation will improve by using better differentiation algorithms (e.g., [25] and [26]).

## Authors’contributions

M.T.A. conceived the project, did the analysis and numerical calculations. All authors analyzed the results. M.T.A. wrote the manuscript with the help of J.X.V-H.

## Acknowledgements

We thank the financial support of CONACyT, México. JXVH acknowledges support from PAPIIT (UNAM) grant IA101215.

## IV. METHODS

A Mathematica notebook to replicate and apply the results of the paper to other time-series is available as Supplementary Material.

### Data

Infection data for measles was obtained from International Infectious Disease Data Archive at McMaster University*. Data for dengue and Zika was obtained from the supplementary information of [20] and [14], respectively. The proportion of infected population was obtained by dividing the reported number of cases *I*(*t_k_*) over the total population provided by a census. The interpolation to obtain *y*(*t*) was done using fourth order splines.

### Used parameters of the algorithm

The parameters of the algorithm were chosen as *μ* = 1.4.*γ* = 1.1 and *ε* = 10^−3^ for measles, *μ* = 3.4, *,γ* = 5.1 and *ε* = 10^−26^ for dengue and *μ* = 1.4, *,γ* = 1.1 and *ε* = 10^−3^ for zika.

### Prediction

For prediction of *β̂*(*t*) over the interval [*t_n_,t_n_* + *ε*], we used the function `TimeSeriesModelFit` provided in the Mathematica software to build a discrete-time linear model of the contact index. In the case of measles, we used the value *β̂*(*t*) from 1940 to *t_n_* = 1941.18 (sampled at rate 0.0001) as training data. The result was the ARIMA discrete-time model `ARIMAProcess` [8.47663 × 10^−13^, {0.0908973}, 4, {0.784174}, 5.06941 × 10^−19^]. In the case of dengue, the training data *β* ̂(*t*) was from 2000.7 to *t_n_* = 2002.18 with sampling rate 0.0001 years. The result was the ARIMA discrete-time model `ARIMAProcess`[−8.94849 × 10^−11^, {1.09715, −0.102606}, 3, {0.518601}, 9.91838 × 10^−16^].

## Supplementary Information

### S1. OBTAINING AN EFFECTIVE REPRODUCTIVE NUMBER FROM THE PROPOSED INDICATOR

The effective reproductive number characterizes a sufficient condition for the end of the epidemic. Hence, the objective in this section is to derive a sufficient condition for the stability of the infected-free solution *I*(*t*) ≡ 0 in the tvSIR model (2). By the continuous dependence of the the solutions of this model on its parameters, we consider the limit case *ε* = 0 only.

Since the differential equation for *M* = *S* + *I* is linear

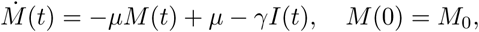

we can compute its solution as

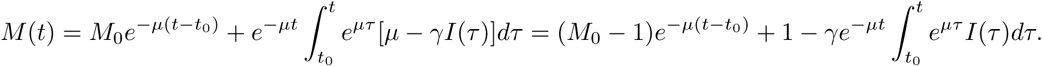

Then, since *S* = *M* − *I* we obtain

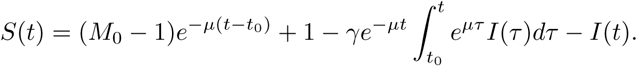

Substituting this expression in (2) means we can rewrite the differential equation for the infected population as

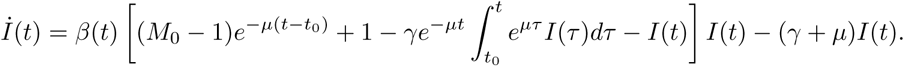

Since *μ* > 0, the term (*M*_0_ − 1)*e*^−*μ*(*t*-*t_0_*)^ → 0 as *t* → ∞ exponentially fast. By definition of stability, only the large time behavior of the system is considered, and hence we obtain the approximation

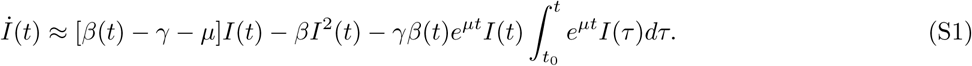

A sufficient condition for stability of the solution *I*(*t*) ≡ 0 is: *I* > 0 implies *İ* < 0. The condition *İ* < 0 in (S1) is equivalent to

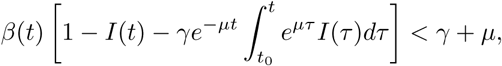

which can be rewritten as R(*t*) < 1 with

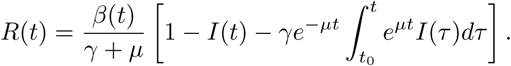

Replacing *β* by *β ̂* obtained in (1) leads to equation (3). Furthermore, since *β ̂*(*t*) → *β* (*t*) exponentially fast, then *R̂*(*t*) → *R*(*t*) exponentially fast also.

**Remark 1.**. A less conservative condition for stability could be obtained by applying the theory of Lyapunov stability (e.g., Lyapunov functionals) to the Volterra differential equation (S1). The analysis here is equivalent to using the Lyapunov function 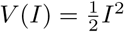 to characterize stability.

**FIG. S1:**
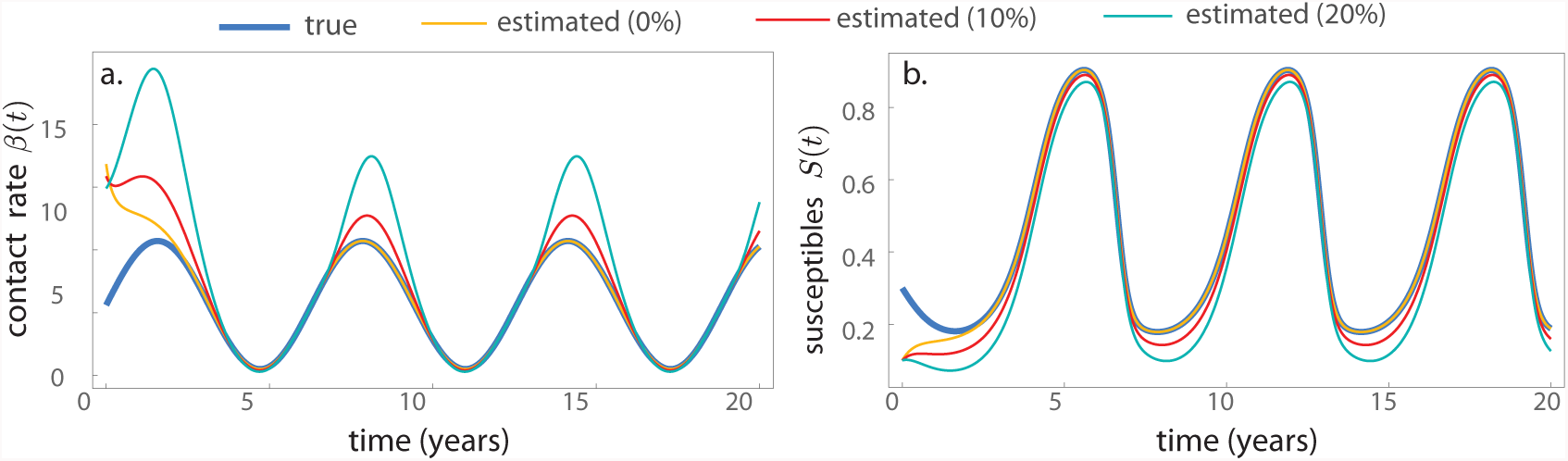
Simulation results of the proposed algorithm for the SIR model. Comparison of the performance of (1) on the SIR model (2) using the exact parameters *γ* = 0.5, *μ* = 1.5, *ε* = 10^−2^, and introducing a 10% error (i.e. (1) uses the parameters 0.9*μ*, 1.1*γ*, 0.9*ε* instead of *γ*, *μ*, *ε*) and 20% error (i.e. 0.8*μ*, 1.2*γ*, 0.8*ε*) in these parameters.

## Footnotes

* https://ms.mcmaster.ca/bolker/measdata.html

